# Body-part specificity for learning of multiple prior distributions in human coincidence timing

**DOI:** 10.1101/2023.10.20.563262

**Authors:** Yoshiki Matsumura, Neil W. Roach, James Heron, Makoto Miyazaki

## Abstract

During timing tasks, the brain learns the statistical distribution of target intervals and integrates this prior knowledge with sensory inputs to optimise task performance. Daily events can have different temporal statistics (e.g. fastball/slowball in baseball batting), making it important to learn and retain multiple priors. However, the rules governing this process are not yet understood. Here, we demonstrate that the learning of multiple prior distributions in a coincidence timing task is characterised by body-part specificity. In our experiments, two prior distributions (short and long intervals) were imposed on participants. When using only one body part for timing responses, regardless of the priors, participants learned a single prior by generalising over the two distributions. However, when the two priors were assigned to different body parts, participants concurrently learned the two independent priors. Moreover, body-part specific prior acquisition was faster when the priors were assigned to anatomically distant body parts (e.g. hand/foot) than when they were assigned to close body parts (e.g. index/middle fingers). This suggests that the body-part specific learning of priors is organised according to somatotopy.

## Introduction

Sensory signals are inherently variable, but Bayesian estimation ^1,2^ can minimise the impact of sensory noise in sensorimotor tasks (e.g. baseball batting in daily tasks). Bayesian estimation involves learning the statistical distribution of a target (e.g. ball speed) and integrating this prior with sensory signals. Psychophysical studies have shown that people behave as predicted by the Bayesian estimation model in various sensorimotor tasks such as reaching ^1^, force matching ^3^, and timing ^4,5^. In most previous studies, a single prior distribution was imposed on the participants within certain task sessions. In daily tasks, however, multiple events occur (e.g. fastball and slowball), and each event can have its own unique statistics. Successful Bayesian estimation in real environments relies on the ability to learn multiple prior distributions.

Recently, Roach et al ^6^ demonstrated that when exposed to two different prior distributions (short and long durations) during a timing task, participants first learned a single prior distribution by generalising over the two distributions (‘generalisation’). Then, after approximately 1000 trials, they eventually learned the two independent priors. Moreover, Roach et al showed that when the two priors were assigned to two different types of motor responses (keypress/vocalisation), participants concurrently learned the two independent priors (‘motor specificity’) within 140 trials. Roach et al. proposed the supplementary motor area (SMA) as a possible neural basis for motor specificity since neurons in the SMA exhibit both time-interval tuning ^7^ and action selectivity ^8^.

In the current study, we hypothesised that when two prior distributions are assigned to two different body parts to generate timing responses, participants may concurrently learn two independent prior distributions, even though the type of motor responses is identical (keypress). This hypothesis has a possible neurophysiological basis—the SMA, which is referred to as a possible neural basis of motor specificity ^6^, also has somatotopic (body-part specific) activity corresponding to the motor outputs ^9,10^. In addition, psychophysical studies suggest that the brain has multiple timers, each of which is associated with different motor effectors (i.e. body parts) ^11^. The body-part specific brain structure and/or function may enable individuals to concurrently learn multiple prior distributions. To test the hypothesis for ‘body-part specificity’, we conducted psychophysical experiments using a coincidence timing task.

## Results

Forty individuals participated in Experiments 1–5 (eight participants per experiment; no overlap among the experiments). They performed coincidence timing tasks (Fig 1a) in which three sequential visual stimuli (S1, S2, and S3) were presented on the right or left of a fixation point. The time interval (*T*_*S*_) between S1 and S2 was equal to that between S2 and S3 in each trial. Based on the *T*_*S*_ from S1 to S2, participants attempted to press a key to coincide with the onset of S3. *T*_*S*_ was randomly sampled from a short (424–988 ms; mean [*μ*_prior_] = 706 ms) or long (1129–1694 ms; *μ*_prior_ = 1412 ms) prior distribution (Fig 1b). Short and long priors were assigned to the left or right stimuli (Fig 1c) in Experiments 1–4 or upper or lower stimuli in Experiment 5. Each participant completed 640 trials of the coincidence timing task (320 trials per prior). The time interval between the onset of S2 and response (*T*_*R*_) (Fig 1a) was used for the analyses.

**Fig 1.**
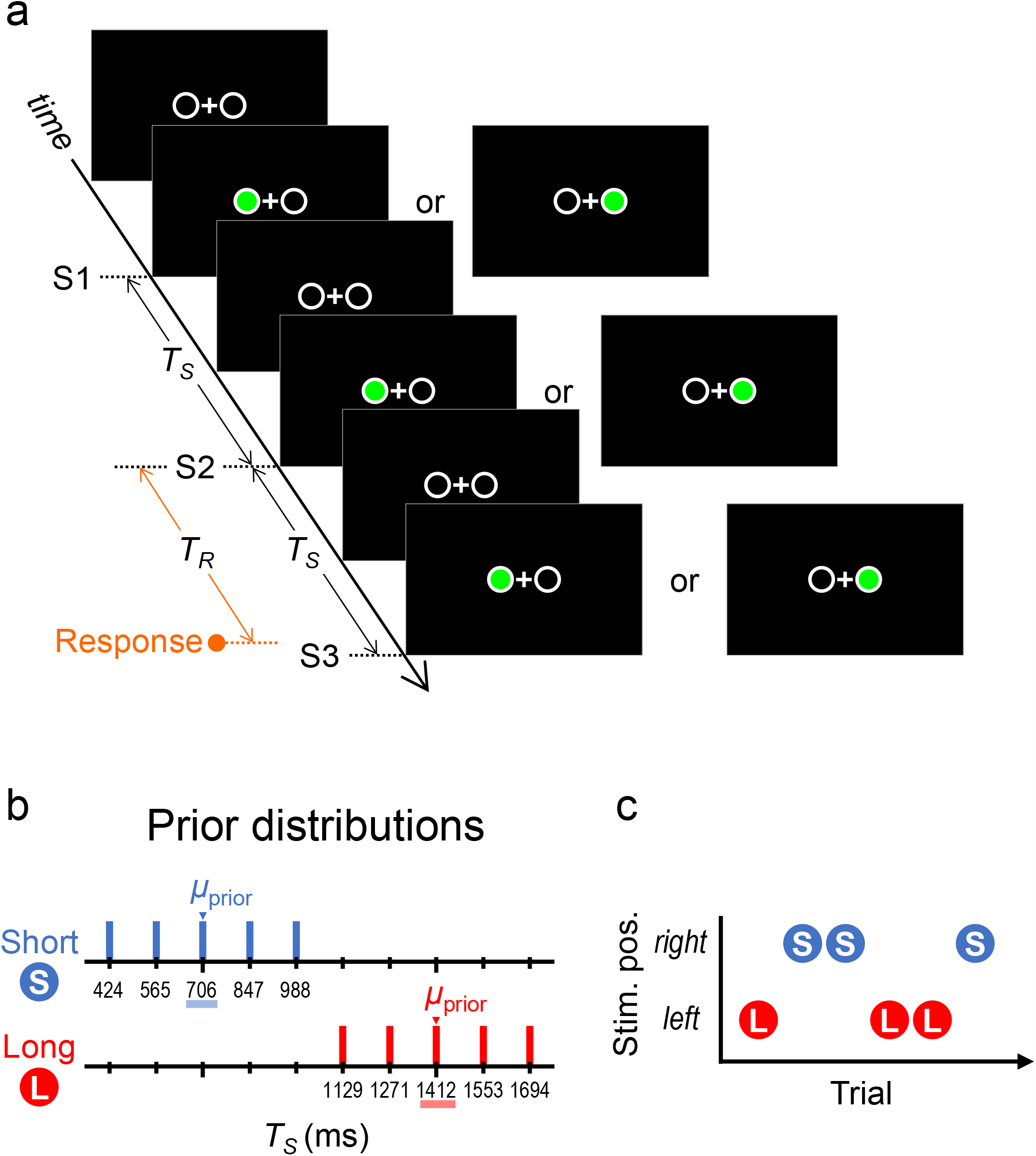
Stimuli and task. **a**: Three sequential stimuli (S1→S2→S3) were presented on the right or left side of the fixation point. The stimulus time interval (*T*_*S*_) between S1 and S2 and that between S2 and S3 were identical in each trial. Based on the *T*_*S*_ from S1 to S2, participants attempted to press a key to coincide with the onset of S3. The time interval from the onset of S2 to that of the response was measured as the response time interval (*T*_*R*_). **b**: *T*_*S*_ was randomly sampled from one of two prior distributions: the short (424, 565, 706, 847, and 988 ms) or long (1129, 1271, 1412, 1553, and 1694 ms) prior. **c**: The short and long priors were assigned to the left- or right-sided stimuli.

### Theoretical predictions

Figs 2a–d show two theoretical predictions for 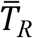 (mean *T*_*R*_ among trials) as a function of *T*_*S*_ (for details, see Theoretical Predictions in Materials and Methods). When Bayesian estimation operates, 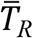 should be biased to the mean of the prior distribution (*μ*_prior_), which is the so called ‘central tendency ^12-14^’. In addition, the bias should be greater with a longer *T*_*S*_ due to scalar variability ^15,16^ (i.e. sensory variability [*σ*_sensed_] is greater under longer *T*_*S*_).

**Fig 2.**
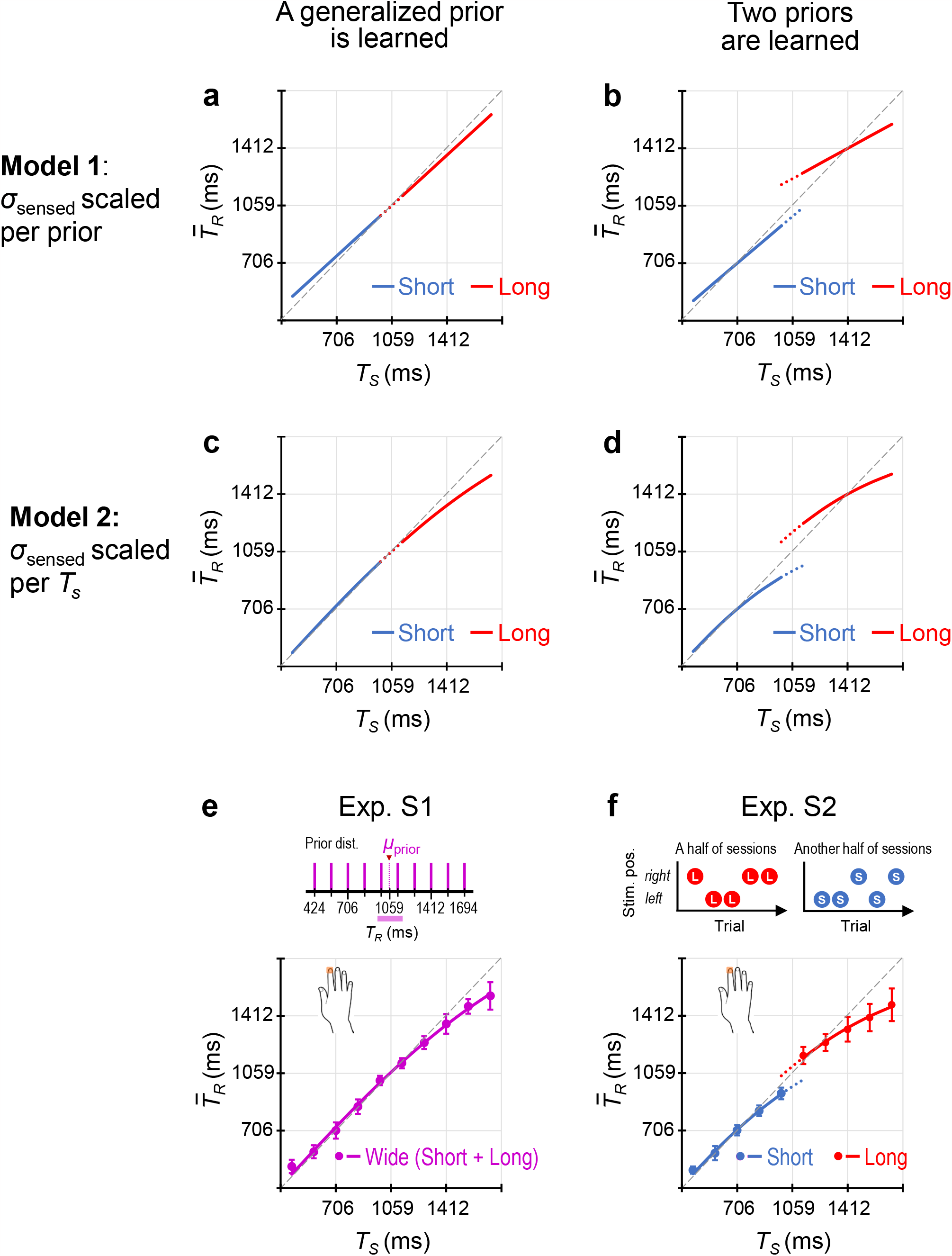
Theoretical predictions by the Bayesian estimation models. In Model 1 (**a, b**), the magnitude of variability in the sensory input of *T*_*S*_ (*σ*_sensed_) is scaled per prior. The mean response time intervals among trials 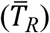 as a function of stimulus time interval (*T*_*S*_) in the case that participants learned a single prior generalised over the two distributions (**a**) and in the case that participants concurrently learned the short and long priors (**b**). In Model 2 (**c, d**), *σ*_sensed_ is scaled per *T*_*S*_. 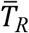 as a function of *T*_*S*_ in the case that participants learned a single generalised prior (**c**) and in the case that participants concurrently learned the short and long priors (**d**). The 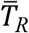 values across the participants (mean ± standard deviation [SD]) as a function of *T*_*S*_ in Supplementary Experiments S1 (**e**) and S2 (**f**), which were calculated using data in the last quarter of trials. In Experiment S1, participants were presented with a single wide prior distribution made by combining the short and long priors. In Experiment S2, participants were presented with only one of the two priors during half of the sessions, and another prior during the other half. The 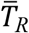 values across the participants supported Model 2.

In Model 1 (Fig 2a, b), *σ*_sensed_ is scaled per prior (i.e. *σ*_sensed_ is constant within each prior), which is explained by a linear equation (Eq. 2 in Theoretical predictions in Materials and Methods) based on previous studies ^5,12-14^. Under this linear model, the central tendency appears as a line with slope < 1 in the 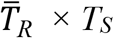 functions. If participants learned a single generalised prior, the lines for the two priors overlap (Fig 2a). If participants concurrently learned the short and long priors, two independent lines appear, and the slope should be lower for the long prior than for the short prior (Fig 2b).

In Model 2 (Fig 2c, d), *σ*_sensed_ scales per *T*_*S*_, which is expressed by a nonlinear equation (Eq. 4). Under this model, the bias of 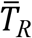 to *μ*_prior_ should be gradually greater with the longer *T*_*S*_. Consequently, the 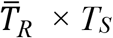 functions exhibit curves with gentler gradients in the longer *T*_*S*_. If participants learned a generalised single prior, the curves for the short and long priors overlap (Fig 2c). If participants concurrently learned the short and long priors, two independent curves appear (Fig 2d).

### Experimental verification of theoretical predictions

To verify which model best describes performance on the coincidence timing task, we conducted two supplementary experiments (for details, see Supplementary Methods and Results, and Supplementary Fig S1). In Experiment S1 (Fig 2e), participants (*n* = 8) were presented with a single wide prior distribution created by combining the short and long priors. In Experiment S2 (Fig 2f), participants (*n* = 8) were presented with the short prior during half of the sessions and the long prior during the other half. The results supported Model 2, which was especially evident in Experiment S1 using a wider range of priors than those used in previous studies ^5,12-14^.

To evaluate how participants learned the two priors, we calculated 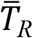 at *T*_*S*_ = 1059 ms (mean of the two priors) on fitted curves [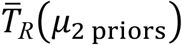, Fig 3a]. If participants learned a single generalised prior, there should be no difference in 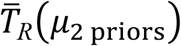 between the priors. However, if participants concurrently learned the two independent priors, 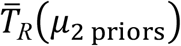 should be greater for the long prior than for the short prior.

**Fig 3.**
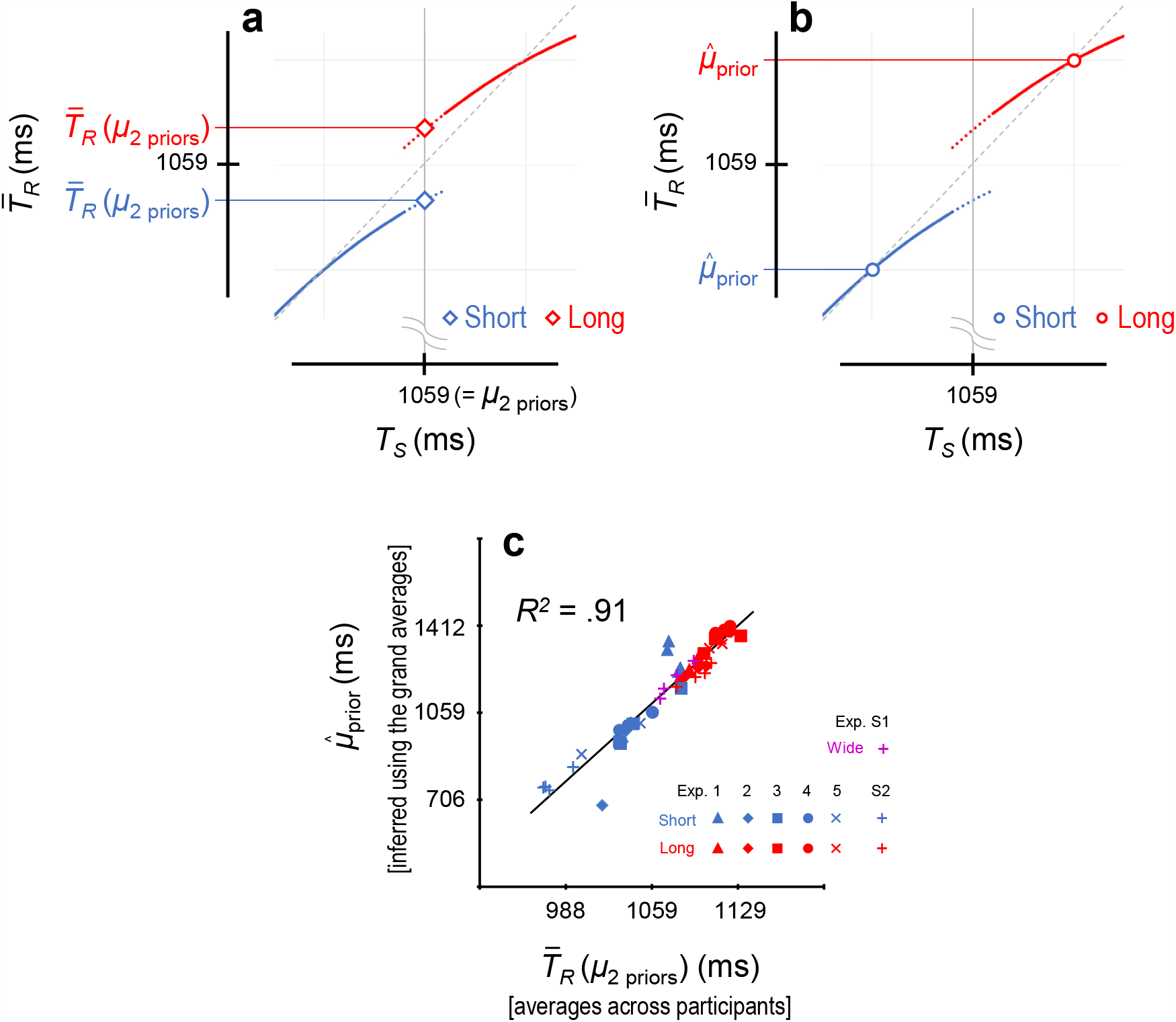
Evaluations of how participants learned the two priors. 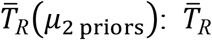 at *T*_*S*_ = 1059 ms on the fitted curves (**a**). 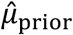: mean of the acquired prior (**b**). 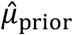 can be inferred from the point that the 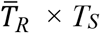 curve intersects the unity line. 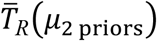 values were calculated for each participant, whereas the 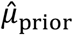 values were calculated using the grand-averaged 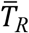 values (means across participants). Correlation between the 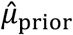 and 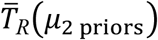 values (**c**). Each plot represents the 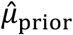and 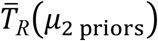 values calculated for each prior per trial-bin in each experiment (for the 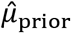 and 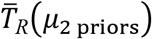 values of Experiments S1 and S2, see Supplementary Fig S1).

In theory, the mean of the acquired prior 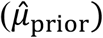 can be inferred from the point that the 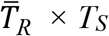 curve intersects the unity line (Fig 3b). In practice, however, the estimation of this intersection was highly sensitive to variability in individuals’ response patterns (for details, see Analyses in Methods). Therefore, we calculated the 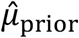 values using the curves fitted to the grand-averaged 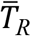 values (i.e. means across participants), in which the individuals’ idiosyncratic responses were cancelled through averaging. As shown in Fig 3c, 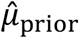 values significantly correlated with 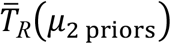 values across participants (*R*^2^ = 0.91).

To facilitate comparison with earlier studies ^12-14^, we also evaluated whether participants concurrently learned the short and long priors using the regression index (1 − slope) that is based on Model 1 (for details, see Analyses in Materials and Methods). If participants concurrently learned the two priors, the regression indices should be larger than zero (i.e. slope < 1) for both priors, and the index should be greater for the long prior than for the short prior.

### Experiment 1: timing using a single body part (index finger)

Participants (*n* = 8) performed the coincidence timing task using only the dominant index finger, regardless of the stimulus locations (i.e. priors) (Fig 4).

**Fig 4.**
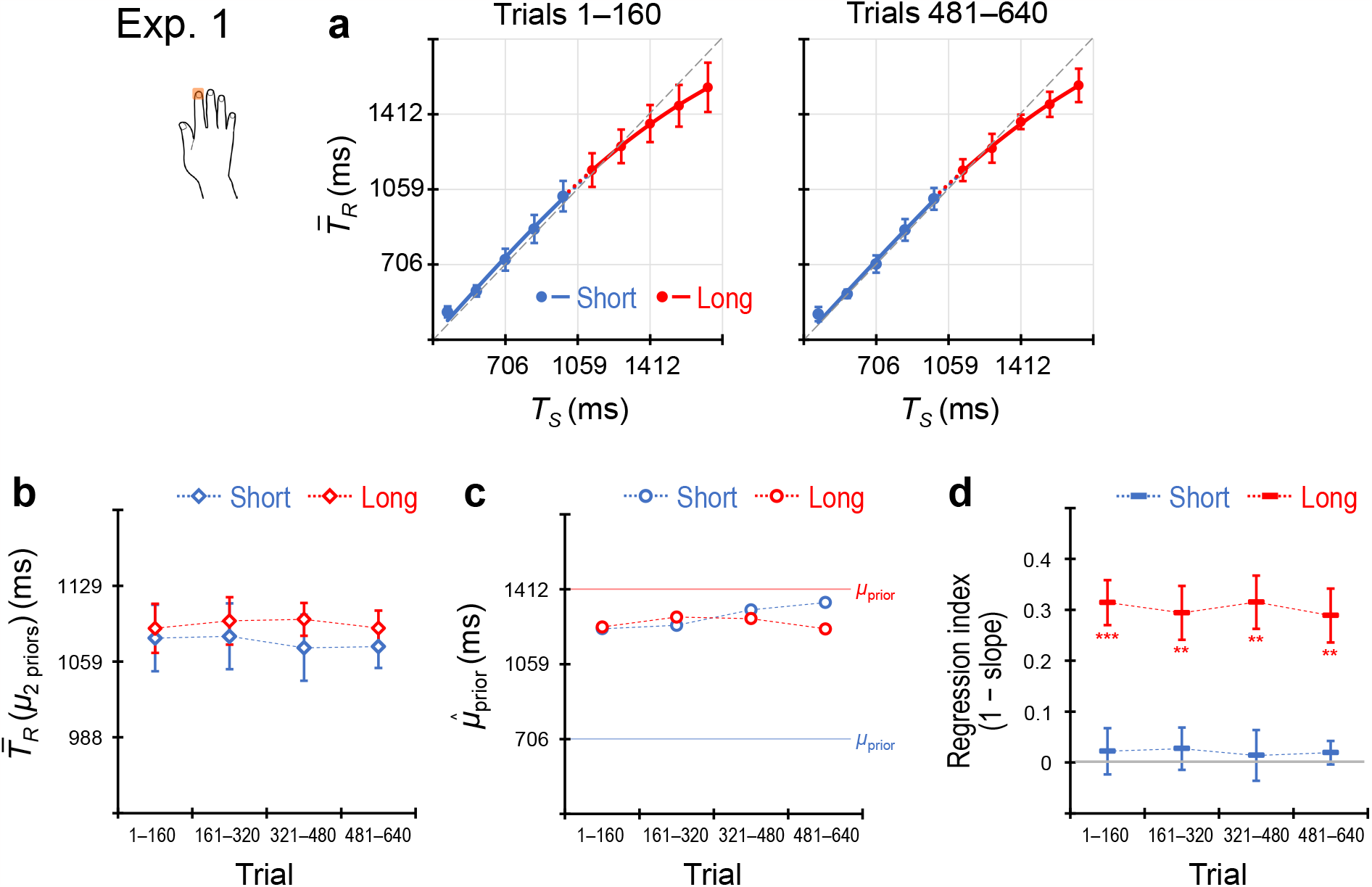
Results of Experiment 1. Participants (*n* = 8) performed the coincidence timing task using only the dominant index finger regardless of the priors. 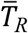 values across participants (mean ± SD) as a function of *T*_*S*_ for trials 1–160 and 481–640 (**a**). 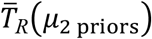 values across the participants (mean ± SEM) for the short and long priors, calculated per 160 trials (80 trials/prior) (**b**). 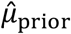values inferred using the grand-averaged 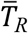 values for the short and long priors, calculated per 160 trials (**c**). Regression indices across participants (mean ± SEM) for the short and long priors, calculated per 160 trials (**d**). ** *p*_*cor*_ < .01, *** *p*_*cor*_ < .001.

Fig 4a shows 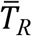 values across participants (mean ± standard deviation [SD]) as a function of *T*_*S*_ for trials 1–160 and trials 481–640 (i.e. trials 1–80 and trials 241– 320 per prior). 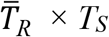 curves for the short and long priors overlapped in both trial bins. Fig 4b shows 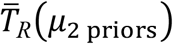 values across participants (mean ± standard error of the mean [SEM]) in each successive 160 trial bins (80 trials/prior). No significant differences between the priors were found (*ps*_*cor*_ ≥ .27, corrected by the Holm method, *ts*(7) ≤ 1.35, Cohen’s *ds* ≤ .48), indicating that participants learned a single prior by generalising over the two distributions. This inference is supported by the 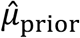 values (Fig 4c), which did not differ systematically between the short and long priors.

Fig 4d shows regression indices across the participants (mean ± SEM), which also did not indicate concurrent learning of the two independent priors. The indices were not significantly greater than zero in all trial bins for the short prior (*ps*_*cor*_ ≥ .39, *t*_7_ ≤ .84, *d* ≤ .30), although those were significantly greater than zero for the long prior (*ps*_*cor*_ ≤ .0026, *ts*(7) ≥ 5.45, *ds* ≥ 1.93) and the difference between the long and short priors remained significant in all trial bins (*ps*_*cor*_ ≤ .0027, *ts*(7) ≥ 3.97, *ds* ≥ 1.40).

### Experiment 2: timing using two body parts (index vs middle fingers)

Next, participants (*n* = 8) performed the coincidence timing task selectively using either the index or middle fingers of the dominant hand according to the stimulus locations (i.e. priors) (Fig 5).

**Fig 5.**
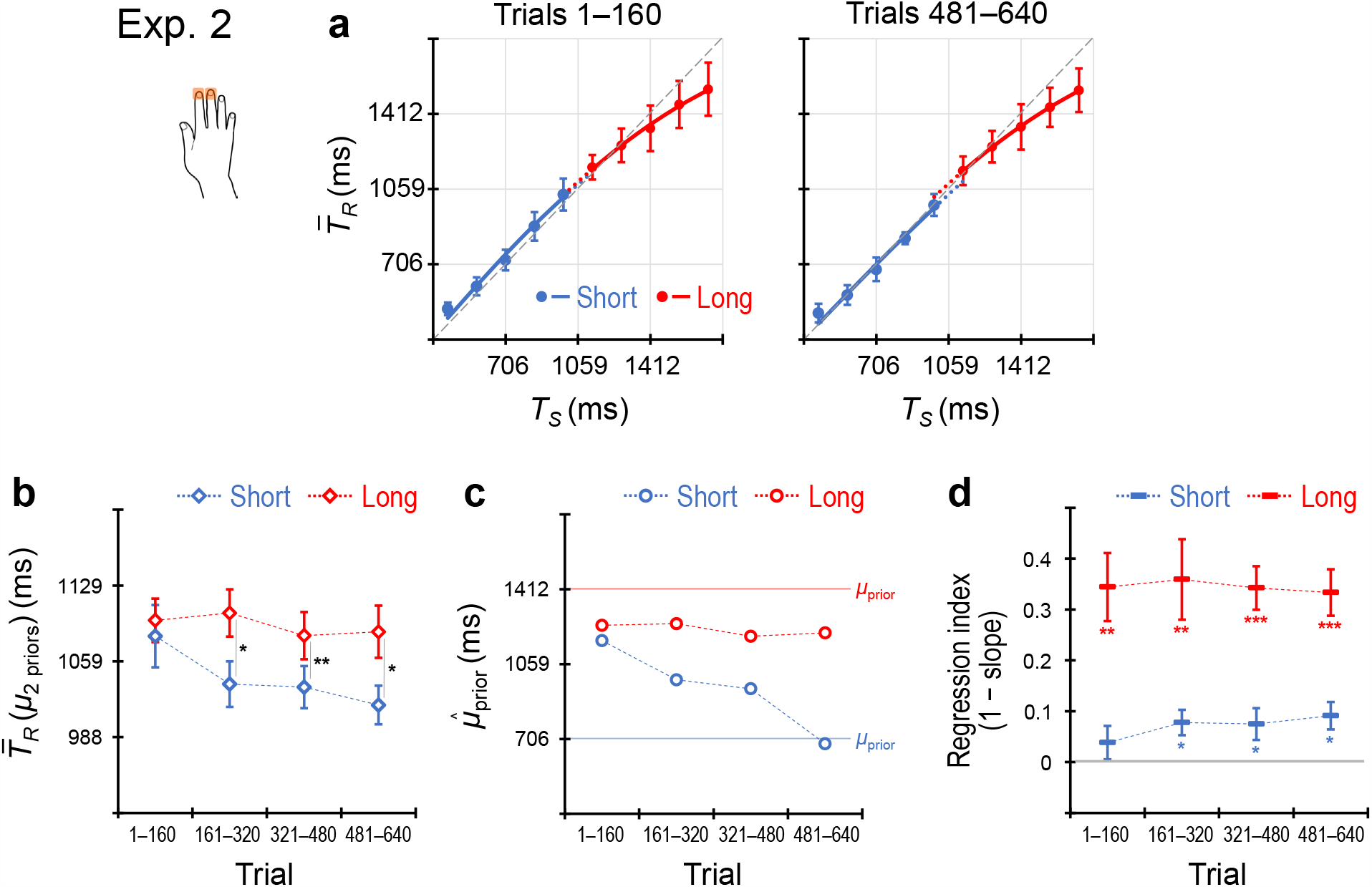
Results of Experiment 2. Participants (*n* = 8) performed the coincidence timing task selectively using the index or middle fingers according to the priors. 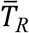 values across the participants as a function of *T*_*S*_ for trials 1–160 and 481–640 (**a**). 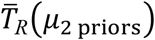 values across the participants for the short and long priors (**b**). 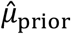values inferred using the grand-averaged 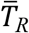values for the short and long priors (**c**). Regression indices across the participants for the short and long priors (**d**). * *p*_*cor*_ < .05, ** *p*_*cor*_ < .01, *** *p*_*cor*_ < .001.

As shown in Fig 5a, 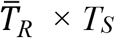 curves for the two priors overlapped in trials 1– 160 but diverged in later trials 481–640. In trials 1–160, 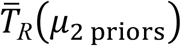 values showed no difference between the two priors (*p*_*cor*_ = .26, *t*(7) = .68, *d* = .24) (Fig 5b). In trials 161–320 and later, however, the 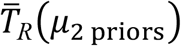 values were significantly greater for the long prior than for the short prior (*ps*_*cor*_ ≤ .025, *ts*(7) ≥ 2.84, *ds* ≥ 1.00). A similar profile is also evident in the 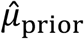 values (Fig 5c), indicating that participants learned a single generalised prior in the early trials before subsequently differentiating the two independent priors.

Regression indices supported the results of the 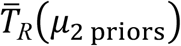 values (Fig 5d). For the short prior, the regression index did not differ significantly from zero in trials 1– 160 (*p*_*cor*_ = .14, *ts*(7) = 1.16, *d* =. 41) but did in later trial bins (*ps*_*cor*_ ≤ .026, *ts*(7) ≥ 2.40, *ds* ≥ .85). The indices for the long prior were significantly greater than zero (*ps*_*cor*_ ≤ .0067, *ts*(7) ≥ 4.54, *ds* ≥ 1.61)), and the difference between the long and short priors remained significant in all trial bins (*ps*_*cor*_ ≤ .0024, *ts*(7) ≥ 4.06, *ds* ≥ 1.43).

The results in Experiment 2 supported our hypothesis of body-part specificity. In the subsequent experiments, we investigated the generality of body-part specificity by using other combinations of body parts.

### Experiment 3: timing using two body parts (right vs left hands)

Participants (*n* = 8) next performed the coincidence timing task selectively using their right or left index fingers according to the stimulus locations (i.e. priors) (Fig 6).

**Fig 6.**
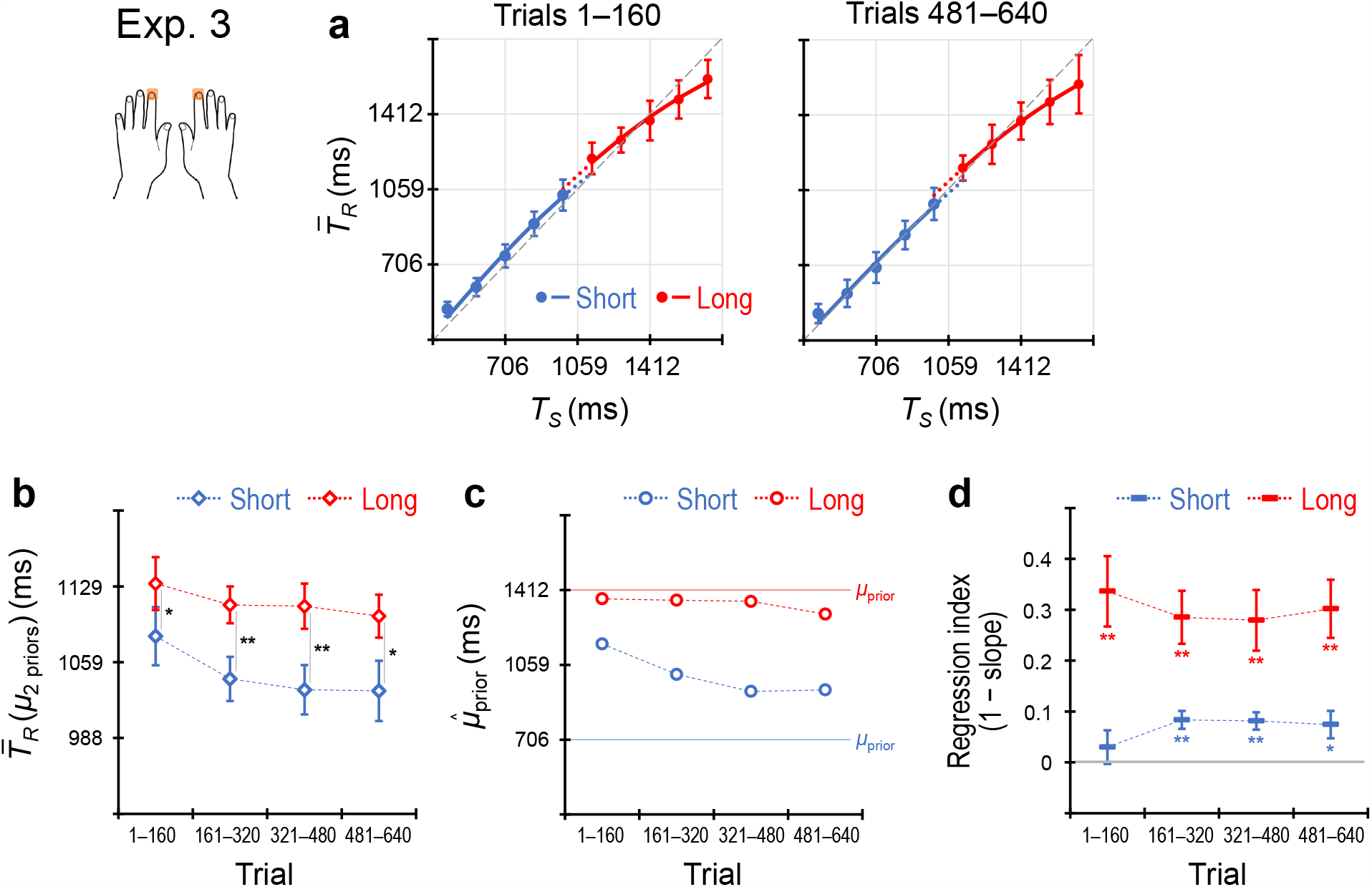
Results of Experiment 3. Participants (*n* = 8) performed the coincidence timing task selectively using the right or left index according to the priors. 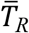 values across the participants as a function of *T*_*S*_ for trials 1–160 and 481–640 (**a**). 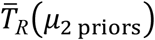 values across the participants for the short and long priors (**b**). 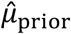values inferred using the grand-averaged 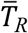values for the short and long priors (**c**). Regression indices across the participants for the short and long priors (**d**). * *p*_*cor*_ < .05, ** *p*_*cor*_ < .01.

In this instance, 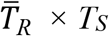 curves diverged between the two priors in both trials 1–160 and 481–640 (Fig 6a) and 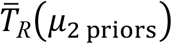 values across the participants were significantly greater for the long prior than for the short prior in all trials (*ps*_*cor*_ ≤ .026, *ts*(7) ≥ 2.33, *ds* ≥ .82) (Fig 6b). The 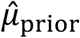 values show similar profiles to the 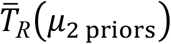 values (Fig 6c). These results suggest that the participants concurrently learned the two independent priors from the early trials.

As shown in Fig 6d, the regression index for the short prior exhibited no significant difference from zero in trials 1–160 (*p*_*cor*_ = .20, *t*(7) = .90, *d* = .32), although the indices were significantly greater than zero in trials 161–320 and later (*ps*_*cor*_ ≤ .029, *ts*(7) ≥ 2.74 *ds* ≥ .97). The indices for the long prior were significantly greater than zero in all trial bins (*ps*_*cor*_ ≤ .0055, *ts*(7) ≥ 4.68, *ds* ≥ 1.66) and greater than those for the short prior in all trial bins (*ps*_*cor*_ ≤ .013, *ts*(7) ≥ 3.43, *ds* ≥ 1.21).

Thus, the regression indices did not indicate central tendency for the short prior in trials 1–160, although did for the long prior. This did not support the concurrent learning of the two independent priors in the early trials, which was not consistent with the results of the 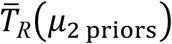 values. Notably, the 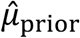 value for the short time prior in trials 1–160 was 1158.1 ms (Supplementary Table S2), which was greater than the short prior (424–988 ms) and was included in the long prior (1129–1694 ms).

According to the Bayesian estimation model, 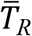 should be biased to 1158.1 ms. In this case, the regression index should be 0 (i.e. slope = 1.0), which is computed from the 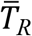 values as a function of *T*_*S*_ with substituting 1158.1 ms, 365.3 ms (Supplementary Table S3), and 0.194 (Supplementary Table S5) (or 1158.1 ms, 256.9 ms [Supplementary Table S4], and 0.136 [Supplementary Table S6]) into *μ*_prior,_ *σ*_prior_, and *w* in Eq. 4, respectively. This is consistent with the experiment result. Thus, Bayesian estimation does not appear as the ‘central’ tendency bias when 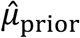 is largely biased towards the other prior.

### Experiment 4: timing using two body parts (contralateral hand vs foot)

Participants (*n* = 8) performed the coincidence timing task selectively using their right/left index finger or left/right heel according to the stimulus locations (i.e. priors) (Fig 7).

**Fig 7.**
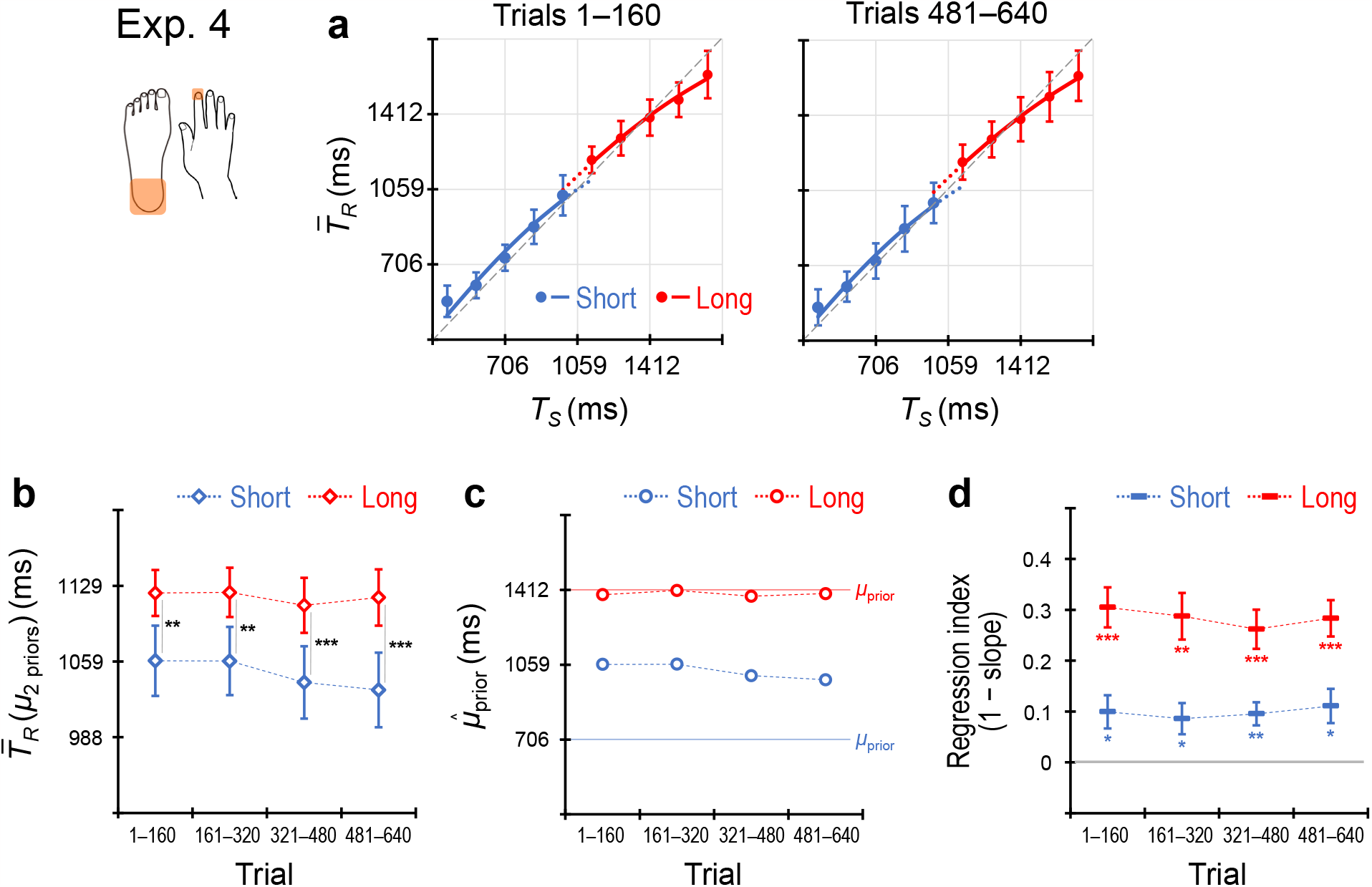
Results of Experiment 4. Participants (*n* = 8) performed the coincidence timing task selectively using the contralateral hand or foot according to the priors. 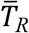 values across the participants as a function of *T*_*S*_ for trials 1–160 and 481–640 (**a**). 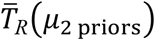 values across the participants for the short and long priors (**b**). 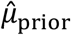values inferred using the grand-averaged 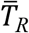 values for the short and long priors (**c**). Regression indices across the participants for the short and long priors (**d**). * *p*_*cor*_ < .05, ** *p*_*cor*_ < .01, *** *p*_*cor*_ < .001.

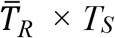 curves for the two priors diverged in both trials 1–160 and 481–640 (Fig 7a). 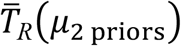 values were significantly greater for the long prior than for the short prior over all trial bins (Fig 7b; *ps*_*cor*_ ≤ .0026, *ts*(7) ≥ 3.99, *ds* ≥ 1.41). The 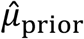 values show a similar profile to the 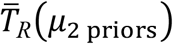 values (Fig 7 c). The results indicate that the participants learned the two independent priors from the early trials.

As shown in Fig 7d, regression indices were significantly greater than zero for the short and long priors in all trial bins (*ps*_*cor*_ ≤ .020, *ts*(7) ≥ 2.80, *ds* ≥ .99). The indices were greater for the long prior than for the short prior in all trial bins (*ps*_*cor*_ ≤ .0021, *ts*(7) ≥ 4.40, *ds* ≥ 1.29). The results were consistent with those of the 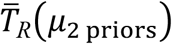 values.

### Experiment 5: timing using two body parts (ipsilateral hand vs foot)

Participants (*n* = 8) performed the coincidence timing task selectively using their dominant index finger or the ipsilateral heel according to the stimulus locations (i.e. priors) (Fig 8). To maintain stimulus–response compatibility, stimuli were presented above or below the fixation point. The participants pressed a key with their index finger when the stimuli were presented above, and they pressed a foot key with their heel when the stimuli were presented below.

**Fig 8.**
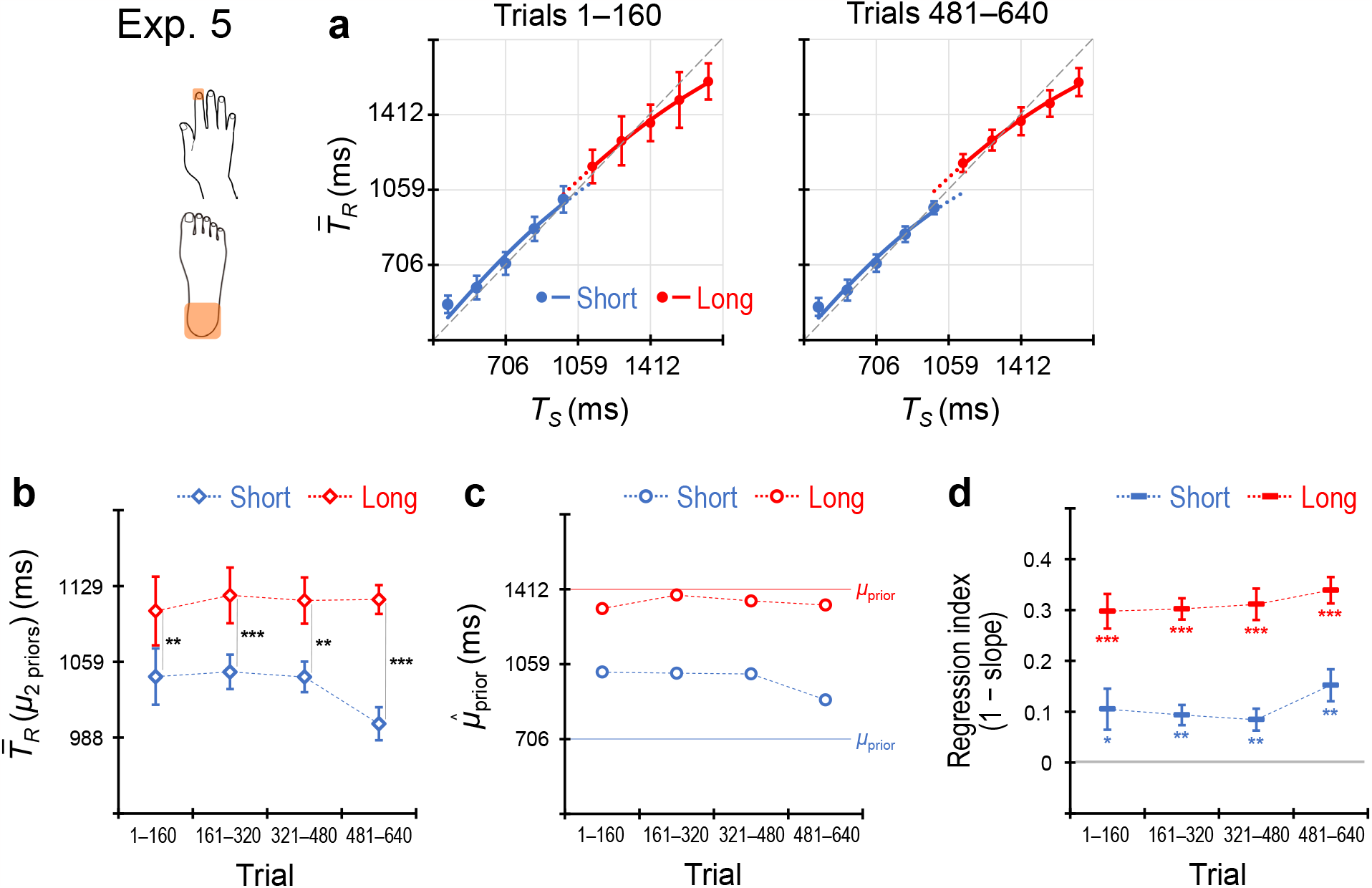
Results of Experiment 5. Participants (*n* = 8) performed the coincidence timing task selectively using the ipsilateral hand or foot according to the priors. 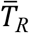 values across the participants as a function of *T*_*S*_ for trials 1–160 and 481–640 (**a**). 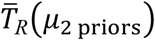 values across the participants for the short and long priors (**b**). 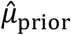 values inferred using the grand-averaged 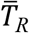 values for the short and long priors (**c**). Regression indices across the participants for the short and long priors (**d**). * *p*_*cor*_ < .05, ** *p*_*cor*_ < .01, *** *p*_*cor*_ < .001.

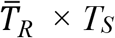 curves for the two priors diverged in both trials 1–160 and 481–640 (Fig 8a), with 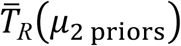 values remaining significantly greater for the long prior than for the short prior over all trial bins (Fig 8b) (*ps*_*cor*_ ≤ .0035, *ts*(7) ≥ 3.94, *ds* ≥ 1.39). The 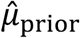 values show a similar profile to the 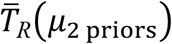 values (Fig 8d). These results indicate that the participants learned the two independent priors from the early trials as in Experiments 3 and 4, even though the two body parts did not cross over the right and left body sides.

Regression indices were significantly greater than zero for the short and long priors in all trial bins (*ps*_*cor*_ ≤ .018, *ts*(7) ≥ 2.60, *ds* ≥ .92) (Fig 8d). The indices were greater for the long prior than for the short prior in all trial bins (*ps*_*cor*_ ≤ .0026, *ts*(7) ≥ 4.00, *ds* ≥ 1.41). The results were consistent with those of the 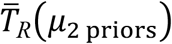 values.

### Comparisons among the experiments

In the results of each experiment, we tested whether participants concurrently learned the short and long priors over time. To directly compare 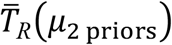 values across experiments, we conducted a three-way analysis of variance with one between-participant factor (5 experiments) and two within-participant factors (2 priors × 4 trial-bins). Significant main effects were found for prior (*F*(1, 35) = 98.73, *p* < .001, 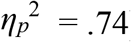) and trial bin (*F*(3, 105) = 4.50, *p* = .0052, 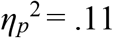), although the main effect of experiment was not significant (*F*(4, 35) = .12, *p* = .98, 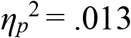). Significant two-way interactions were found between experiment and prior (*F*(4, 35) = 3.83, *p* = .011, 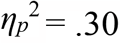) and between prior and trial bin (*F*(3, 105) = 5.98, *p* < .001, 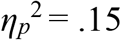). However, the interaction between experiment and trial bin (*F*(12, 105) = .74, *p* = .71, 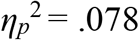) and the three-way interaction of all three factors (*F*(12, 105) = 1.14, *p* = .33, 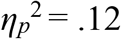) were not significant.

The analyses of simple effects for the interaction between experiment and prior revealed that the effect of prior was non-significant in Experiment 1 (*F*(1, 7) = 1.98, *p* = .20, 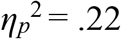) but significant in Experiments 2–5 (*Fs*(1, 7) . 12.50, *ps* ≤ .0095, 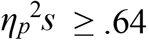). The results further supported that a single generalised prior was learned in Experiment 1 (Fig 4), whereas the two independent priors were generally learned in Experiments 2–5 (Fig 5–8).

The analyses of simple effects for the interaction between prior and trial bin revealed that the effect of trial bin was significant for the short prior (*F*(3, 105) = 7.35, *p* < .001, 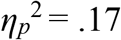) but non-significant for the long prior (*F*(3, 105) = .81, *p* = .49, 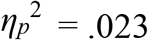). Consistent with the statistical results, 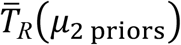 and 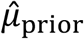 values for the long prior showed little changes among trial bins in all experiments but those for the short prior decreased over trials in some experiments (e.g. Exp. 2, 3, and 5). Further analysis of the simple effect of trial bin for the short prior in each experiment revealed that the effect was significant in Experiments 2, 3, and 5 (*Fs*(3, 21) ≥ 3.14, *ps* ≤ .047, 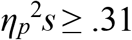) but not in Experiments 1 and 4 (*Fs*(3, 21) ≤ 1.64, *ps* ≥ .21, 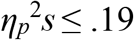).

These results suggest differences in the prior learning processes among the experiments as follows. In Experiment 1, a single generalised prior was acquired. In Experiments 2, 3, and 5, the acquisition of the long prior was completed in the early trials, whereas that of the short prior was attained or developed as the trials progressed. In Experiment 4, the acquisition of the two independent priors was completed in the early trials.

## Discussion

### Body-part specificity for learning of multiple prior distributions

The current results supported our hypothesis of body-part specificity. When participants used only one body part for timing responses, they learned a single prior by generalising over two prior distributions (Exp. 1). However, when participants selectively used two body parts, they concurrently learned the two independent priors (Exp. 2–5). Moreover, the results revealed that body-part specific learning of the priors was more quicky attained when the priors were assigned to anatomically distant body parts (e.g. hand vs foot) than when they were assigned to closer body parts (e.g. index vs middle finger). Thus, body-part specific learning of the priors was regulated in a somatotopic manner.

### Possible neural bases of body-part specificity

We hypothesised body-part specificity based on a previous proposal ^6^ and somatotopy in the SMA ^9,10^. The current results supported the hypothesis and exhibited further consistency with the neurophysiological properties of the SMA. The hand and leg movements are mapped in the relatively rostral and caudal regions in the SMA, respectively ^10^. More than half of the neuronal activity in the SMA is contralateral to the moving hand and foot ^9^. Thus, the body parts that are anatomically distant (or close) are also represented in the anatomically distant (or close) regions in the SMA. Accordingly, we infer the following somatotopic organisation of body-part associated learning of the priors: When the neural representations of the body parts in the SMA are identical and close, the assigned priors are generalised. However, when those in the SMA are distant, the assigned priors are independently acquired.

Moreover, neurons in the dorsal frontal regions including the SMA (pre-SMA and SMA proper) of monkeys exhibited activity consistent with their Bayesian timing behaviour ^17^. In addition, the SMA has neuronal connectivity with the basal ganglia (BG) and cerebellum ^18^. The BG is a core neural basis that encodes time intervals and exhibits neuronal responses that reflect scalar variability ^15^. Narain et al ^19^ showed that the neural circuit model of the cerebellum can learn and represent the prior distribution of time intervals, and the outputs of the circuit replicated the psychophysical observations ^4,5^. It is reasonable to hypothesise that the SMA, BG, and cerebellum constitute a neuronal network to generate Bayesian estimation that can learn multiple priors in timing.

### Preferential acquisition of the long prior

An unexpected feature of our results was that, while participants quickly acquired the long prior, they needed more trials to learn the short prior (Exp. 2, 3, and 5). Similar results were found in Experiment S1 but not in Experiment S2 (see Supplementary Results and Supplementary Fig S1). Therefore, such preferential acquisition of the long prior occurred when the short and long priors were intermixed within a session.

A similar effect was also found in some conditions in experiments by Roach et al ^6^. When participants eventually learned the two priors after excessive trials, it was attained by shifting the 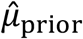 for the short prior away from that for the generalised prior (Fig 4 in Roach et al). Meanwhile, when the two independent priors were learned by adding vocalisation (i.e. motor specificity), the 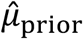 for the long prior was shifted compared to the generalised prior (Fig 5c in Roach et al).

Although the mechanism for the preferential acquisition of the long prior is unclear at this stage, this phenomenon likely appears when participants used only the keypress-type motor responses. In contrast, the opposite effect was observed when participants also used vocalisation. This contrast may reflect some difference in the neuronal or psychological process between body-part specificity and motor specificity.

### Implication of the current results and future perspectives

The body-part specificity found in the present study does not necessarily imply that prior distributions are represented or implemented within motor processing.

Psychophysical ^14^ and psychophysiological ^20^ studies indicated that the prior distribution affects the perceptual process during timing tasks. There is growing evidence that motor responses or programs affect various types of time perception ^21-25^. Thus, it is also plausible that motor responses affect the Bayesian learning of perceptual timing.

Notably, a meta-analysis of neuroimaging studies showed that the SMA was consistently activated across various motor and perceptual timing tasks ^26^. We proposed the SMA as a possible neural basis of body-part specificity. If it is true, the current and previous results are consistent with each other. In future studies, neuroimaging or neurophysiological experiments are necessary to verify this hypothesis.

There is not only neuronal plausibility but also functional significance in motor-perception coupling in the Bayesian learning of timing. During sensorimotor tasks, the variety of motor responses is generally greater when using multiple body parts than when using only a single body part. It should be rational to increase the variety of the perceptual strategy according to the increase in the variety of the motor responses. Inversely, when the variety of the motor response is limited, it would be reasonable for efficient utilisation of a finite perceptual resource to narrow down variety in the perceptual strategy.

In the current study, we focused on the means of the prior distributions acquired in the brain 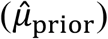 to evaluate whether participants learned the two independent priors. It remains unclear as to how participants learn the variability (*σ*_prior_) of the prior distributions. The *σ*_prior_ acquired in the brain 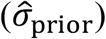 and Weber fraction (*w*) are unspecified by the fitted curves (*cf*. Supplementary Tables S3–S6). Therefore, we cannot directly discuss the 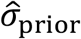 values. Meanwhile, the 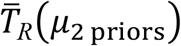 and 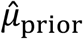 values linearly correlated across the experiments (Fig 3c), on which the values for the wide prior (Exp. S1) plotted with little deviation (purple crosses). This suggests that there was little difference in the 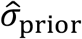 and *w* values between Experiment S1 and other experiments (see Analyses and Eq. 5), although the wide prior had a wider *σ*_prior_ than the short and long priors. It was suggested that the learning of *μ*_prior_ is attained by a smaller number of trials ^27^, whereas that of *σ*_prior_ needs a greater number of trials ^5^. The participants might first acquire a generalised wide 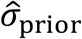 and not be able to fully learn the two independent narrower 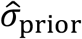 within 640 trials, even when they learned the 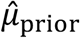 of the two independent priors. Previous studies also measured the *w* values by using time-interval judgment tasks ^12,13^. This methodology enables to directly evaluate both 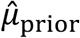 and 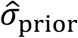 values in future studies.

Whether body-part specificity also operates in nontemporal tasks remains unclear as well. Notably, as a ‘side effect’, the association between the body parts and prior distributions was found in the judgement of visual motion directions ^28^, although the causal relationship between the motor responses and priors was opposite to that in our tasks. In this study, participants triggered the motion stimulus (upward/downward) by key pressing of the right or left index finger. Then, they judged the motion direction. The right and left keypresses were associated with either of two prior distributions (upward/downward). Consequently, their judgments were biased to the means of the priors. The ecological validity of this effect was unclear, since such a causality from motor responses to visual statistics is not easily found in daily environments. However, this finding suggests that the brain is capable of associating the body parts with priors in the nontemporal task. In addition, in a spatial aiming task, dependency on a prior differed according to whether a typical reaching or atypical wrist-rotation setting was used ^29^. Based on the result, the authors proposed that a single prior could be learned differently across different effectors (finger/wrist). This mechanism may also enable to learn multiple priors in the spatial task.

Moreover, it will be instructive to investigate daily human behaviours such as sports ^30-33^. Skilful baseball or cricket players may utilise body-part specificity. For example, to improve the hit rates, a batter may beat time using a foot when a fastball is pitched (i.e. short prior) whereas using a hand when a slowball is pitched (i.e. long prior). In addition, future studies on people with autism can further the current findings. Previous studies demonstrated that individuals with autism or high autistic traits have a disability in learning the prior distribution in time processing ^13,34^. People with autism may also exhibit their specific behaviour in the learning of the multiple prior distributions. Our finding would enhance the application of the Bayesian approach to our daily behaviour.

## Materials and Methods

### Participants

Forty healthy individuals participated in Experiments 1–5. In addition, 16 healthy individuals participated in Supplementary Experiments S1 and S2 (see Supplementary Methods). Eight individuals participated in one of seven experiments (for the profiles of participants in each experiment, see Supplementary Table S1). There was no overlap of participants among the experiments to avoid a possible effect of the priors learned in the previous experiment. All participants were naïve to the purpose of the experiments.

This study was approved by the Ethics Committee of Shizuoka University (15-19). All experiments were performed in accordance with the approved guidelines and regulations. All participants provided written informed consent.

### Stimuli

In a dimly lit sound-shielded room, each participant placed their head on a chin rest and sat at the distance of 87 cm from the monitor (Sony GDM-F500, Japan; 85 Hz). Presentation software (Neurobehavioral Systems, USA) was used for generating the stimuli and recording responses of the participants.

Three sequential stimuli (S1→S2→S3) were presented on the right or left side of the fixation point (Fig 1a) in Experiments 1–4 or the upper or lower side in Experiment 5. The duration of each stimulus was 106 ms. The diameter of the frame circles in which stimuli (green emission) appeared was 1.1° in visual angle, and the distance between the centre of the two circles was 2.2°. The stimulus time interval (*T*_*S*_) between S1 and S2 and that between S2 and S3 were identical within a trial.

For each trial, *T*_*S*_ was randomly sampled from either of two discrete uniform prior distributions (Fig 1b): the short prior (424, 565, 706, 847, and 988 ms; *μ*_prior_ = 706 ms) or the long prior (1129, 1271, 1412, 1553, and 1694 ms; *μ*_prior_ = 1412 ms). The short and long priors were assigned to the left or right stimuli (Fig 1c) in Experiments 1–4 or the upper or lower stimuli in Experiment 5. The combinations between the priors (short/long) and stimulus locations (right/left or upper/lower) were counterbalanced among participants in each experiment. The trial-by-trial order of the priors (i.e. stimulus locations) was randomly determined with the restriction that *T*_*S*_ was not repetitively sampled from the same prior for more than four trials.

### Task

Based on *T*_*S*_ from S1 to S2, participants attempted to press a key to coincide with the onset of S3 (coincidence timing task). For each trial, the time interval from the onset of S2 to that of the motor response was measured as the response time interval (*T*_*R*_) (Fig 1a). The key pressing was conducted using the dominant index finger only (Exp. 1, S1, and S2), dominant index or middle finger (Exp. 2), left or right index finger (Exp. 3), right/left index finger or left/right heel (Exp. 4), or dominant index finger or ipsilateral heel (Exp. 5) according to the stimulus locations (left/right or upper/lower) with stimulus–response compatibility. A single key was used in Experiment 1. Two keys were used in Experiments 2–5. The horizontal distance between the centres of the left and right keys was 1.9 cm in Experiment 2, 11.5 cm in Experiment 3, and that between the contralateral hand and foot keys was 11.5 cm in Experiment 4. The centres of the hand and foot keys were approximately placed at the same horizontal position in Experiment 5.

### Procedure

Each participant completed 640 trials (40 trials/session × 16 sessions) of the task. The interval from the onset of S3 in a trial to that of S1 in the subsequent trial was 4.1 sec. A short beep (0.2 s) was presented 1 s before S1 to alert participants to the beginning of the trial. Participants took a 1-min break after each session and a 5-min break per 4 sessions. When participants reported fatigue or drowsiness, the break was extended.

### Theoretical predictions

According to the Bayesian estimation theory ^1,2^, the brain integrates the sensory input of a target (*X*_sensed_) and the prior distribution about the target to obtain the optimal estimate of the target (*X*_estimated_), as follows:

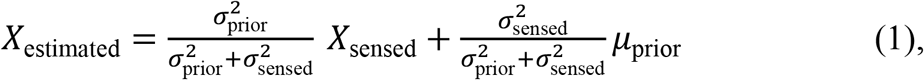

where *σ*_sensed_ denotes the SD of *X*_sensed_ (i.e. degree of sensory variability). *μ*_prior_ and *σ*_prior_ denotes the mean and SD of the prior distribution.

Assuming that there is no bias at the stages of sensory inputs and motor outputs in the coincidence timing task, *X*_sensed_ can be approximated by *T*_*S*_ and the mean among trials of *X*_estimated_ can be approximated by that of 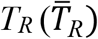. Accordingly, 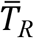 can be expressed as a linear function of *T*_*S*_ as follows:

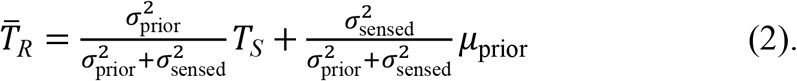

In Eq. 2, we assumed the prior distribution as a Gaussian distribution, although the uniform distributions were used for the priors, based on previous studies ^12-14,35^. In Eq. 2, 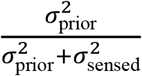 represents the slope of 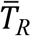 against *T*_*S*_, and 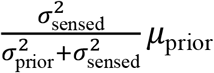 represents the intercept.

Fig 2a and b show the theoretical predictions for the experimental results according to Eq. 2 (Model 1). In each 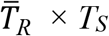 function, the dashed diagonal line expresses the unity line (slope = 1, intercept = 0). In the case that participants used no prior distribution (modelled by *σ*_prior_ = ∞), 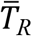 should be plotted on the dashed line. However, when participants learned a prior distribution, the slope should be smaller than 1 (‘central tendency’). The central tendency reflects that the estimate of *T*_*S*_ is biased to *μ*_prior,_ due to Bayesian estimation. The greater central tendencies appear as the smaller slopes.

Fig 2a shows the prediction when participants learned a single prior by generalising over the short and long distributions (‘generalisation’). In this case, the central tendency occurs toward the mean between the short and long priors (*μ*_2 priors =_ 1059 ms in our experiments). Accordingly, 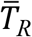 for the two priors should be plotted on a common line with slope < 1. Fig 2b shows the prediction when participants concurrently learned the short and long priors. In this case, the two slopes are smaller than 1. In addition, the slope is smaller for the long prior than for the short prior. This is due to scalar variability ^15,16^. According to scalar variability, greater *σ*_sensed_ arises under the long prior than under the short prior, which causes a greater bias to the *μ*_prior_ (i.e. smaller slope) for the long prior. This model was used in a previous study and was well fitted to the results ^13^.

Fig 2a and b show the predictions with an assumption that *σ*_sensed_ is scaled per prior. That is, *σ*_sensed_ is identical for all *T*_*S*_ within each prior. However, according to the original concept of scalar variability, *σ*_sensed_ should be scaled per *T*_*S*_ as follows:

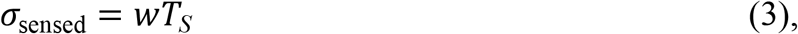

where *w* denotes the Weber fraction. Then, Eq. 2 can be transformed as

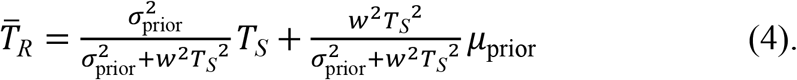

Eq. 4 predicts that the 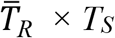 functions exhibit nonlinear profile with a gentler gradient in the longer *T*_*S*_ (Fig 2c and d, Model 2). Fig 2 c shows the prediction when participants learned a single generalised prior. Fig 2d shows the prediction when participants concurrently learned the short and long priors.

As shown in Fig 2e, f, the results of supplementary Experiments S1 and S2 indicated that Model 2 was adequate for explaining those of Experiments 1–5 (for details, see Supplementary Methods and Results and Supplementary Fig S1).

### Analyses

We measured *T*_*R*_ in each trial (Fig 1a). We excluded trials containing any of the following responses from analyses: no key pressing, pressing the opposite key, pressing the key twice or more. We sorted the *T*_*R*_ values for each *T*_*S*_ every 160 trials (80 trials per prior) to calculate the 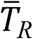 values for each prior. Then, we fitted the 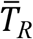 values as a function of *T*_*S*_ by the nonlinear function of Eq. 4 using the least squares method.

As shown in Fig 3a, we calculated 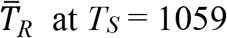 ms (i.e. *μ*_2 priors_) on the fitted curves 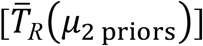. The 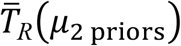 values should have no difference between the short and long priors if participants learned a single prior by generalising over the two distributions. Meanwhile, the 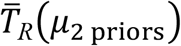 values should be greater for the long prior than for the short prior if participants concurrently learned the two independent priors.

We calculated the 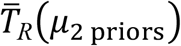 values for each prior per trial bin in each participant. Then, we tested whether the 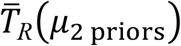 values were greater for the long prior than for the short prior in each experiment, using one-tailed paired *t*-tests corrected by the Holm method (corrected *p*-values [*p*_*cor*_] are shown). Moreover, for the comparisons among the experiments, we conducted a three-way analysis of variance with one between-participant factor (experiment) and two within-participant factors (prior, trial-bin) on the 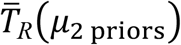 values.

In theory, the mean of the acquired prior 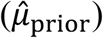 can be inferred from the point that the 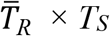 curve intersects the unity line (i.e. the point of 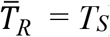) (Fig 3b). Substituting 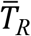 into *T*_*S*_ (or *T*_*S*_ into 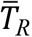), Eq. 4 derives 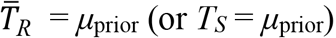. The *μ*_prior_ estimated from the participant’s responses should reflect the 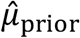. However, the estimation of the intersection is highly sensitive to idiosyncratic responses (e.g. overshoots, undershoots, steeper slopes than unity), leading to implausible 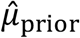 values (e.g. < 0 ms, > 2000 ms) being obtained in 14.4 % of the curve fittings. Therefore, we could not use the 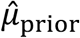 values for the statistical tests.

Instead, we calculated the 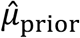 values using the grand-averaged 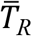 values (mean across participants). There was no implausible value when inferring 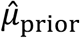 using the curves fitted to the grand-averaged 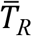 values because idiosyncratic responses for each participant were cancelled by averaging across the participants. As shown in Fig 3c, the 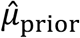 values linearly correlated with the 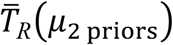 values across participants among the experiments, priors, and trial bins (*R*^*2*^ = 0.91). Thus, the 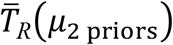 values across participants can be used for explaining the 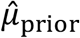 values.

In theory, 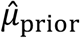 can be expressed as a function of 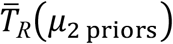 as follows:

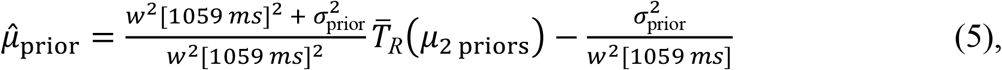

which can be obtained by resolving Eq. 4 after substituting 1059 ms and 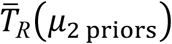 into *T*_*S*_ and 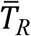, respectively. Assuming that σ_prior_ and *w* are constant, Eq. 5 is a linear function of 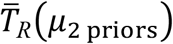. The significant correlation between the 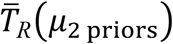 and 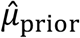 values (Fig 3c) might imply that there was no difference in the *σ*_prior_ and *w* values among the experiments, priors, and trial bins. Alternatively, although there might be a difference in *σ*_prior_ or *w* among them, the effects would be relatively too small to affect the linear correlation between the 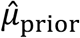 and 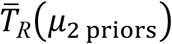 values.

In addition, we evaluated the degree of the central tendency for the short and long priors in Experiments 1–5 and S2 using the regression index ^12,13^, which was calculated by subtracting the slope of the regression line of 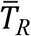 against *T*_*S*_ from one (1 – slope). As shown in the results of Experiment S1 (Fig 2e), the 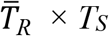 function should be essentially modelled by the nonlinear function as Eq. 4 in the current experiments. However, the linear slope has been widely used for evaluating the central tendency in previous studies ^5,12-14^. There should be no practical problem approximating the 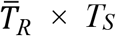 function using the linear function as in Eq. 2 when calculating within short ranges of *T*_*S*_ as done in previous studies (e.g. max – min ≈ 0.5 s). To facilitate a comparison with earlier studies, it is reasonable to evaluate the results also using the regression index.

If participants concurrently learned the two independent priors, the regression indices would be greater than zero for both short and long priors. In addition, the regression index should be greater for the long prior than for short prior due to scalar variability. We tested them per 160 trials (80 trials/prior) in each experiment using the one-tailed paired *t*-test corrected by the Holm method. Notably, the criterion of the regression index > 0 is not always a necessary condition to verify that participants learned the two independent priors when 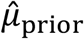 is largely biased to the other prior (see Results of Experiment 3).

## Author Contributions

M.M. conceived and supervised the study. N.W.R. and J.H. helped the experimental design. N.W.R. and M.M. developed the theoretical models. Y.M. set up and conducted the experiments. Y.M. and M.M. analysed data. All authors discussed the results. M.M. wrote the manuscript. N.W.R. and J.H. helped the writing of the manuscript.

## Acknowledgements

The authors thank R. Sato, T. Sato, and J. Ikki for technical assistance with the experiments and N. Enomoto and Y. Sato for administrative assistance with the experiments.

This study was supported by JSPS KAKENHI (grant numbers 22H00502, 22K18263, 19H01087, and 17KK0004)

## Data Availability

The datasets generated and/or analysed during the current study are available from the corresponding authors on reasonable request.

## Code Availability

The underlying code for this study is not publicly available but may be made available to qualified researchers on reasonable request from the corresponding author.

