## Supplementary Information for "Body-part specificity for learning of multiple prior distributions in human coincidence timing"

### Supplementary Methods

#### *Experiments S1 and S2*

#### Participants

Sixteen healthy individuals participated in Supplementary Experiments S1 and S2. Eight individuals participated in one of the two supplementary experiments. There was no overlap of participants among Experiments 1–5 and Supplementary Experiments S1 and S2 (see Supplementary Table S1 for the profiles of the participants).

**Supplementary Table S1. Profiles of the participants**

| Exp. # | *n* | Female/Male | Left/Right-hande | Age [mean ± SD (min–max)], years |
|---|---|---|---|---|
| 1 | 8 | 1/7 | 1/7 | 20.8 ± 1.3 (18–22) |
| 2 | 8 | 3/5 | 1/7 | 22.1 ± 2.6 (19–26) |
| 3 | 8 | 2/6 | 0/8 | 22.0 ± 1.7 (20–25) |
| 4 | 8 | 2/6 | 2/6 | 21.4 ± 1.6 (19–24) |
| 5 | 8 | 3/5 | 2/6 | 21.1 ± 2.4 (19–26) |
| S1 | 8 | 1/7 | 1/7 | 21.5 ± 1.1 (20–23) |
| S2 | 8 | 3/4 | 1/7 | 20.8 ± 1.4 (18–23) |

#### Stimuli, Task, and Procedure

The stimuli, task, and procedure in Experiments S1 and S2 were the same as those in Experiments 1–4 except for the following points. In Experiment S1, $T_S$ for both the right and left stimuli were randomly sampled from the wide prior distribution created by combining the short and long priors (upper part in Fig. 2e) during all sessions. In Experiment S2, $T_S$ for both the right and left stimuli were randomly sampled from either the short or long prior distributions during half of the sessions and another prior during the other half (upper part in Fig. 2f). The participants conducted the coincidence timing task by pressing a key using only the dominant index finger in Experiments S1 and S2 as in Experiment 1.

#### Analyses

There were no differences in $\bar{T}_R$ values across the participants between the right- and left-sided stimuli in both Experiments S1 and S2. Therefore, we averaged the $T_R$ values for the right- and left-sided stimuli to calculate the $\bar{T}_R$ values for each $T_S$, then we calculated the $\bar{T}_R(\mu_{2\ \mathrm{priors}})$ and $\hat{\mu}_{\mathrm{prior}}$ values in Experiments S1 and S2 and regression indices in Experiment S2.

A three-way repeated-measures analysis of variance (ANOVA) (2 stimulus sides × 10 $T_S$ × 4 trial-bins) of $\bar{T}_R$ values in Experiment S1 showed no significant main effect of stimulus side ($F(1, 7) = .16$, $p = .70$, $\eta_p^2 = .022$), although there were significant main effects of $T_S$ ($F(9, 63) = 1616.03$, $p < .001$, $\eta_p^2 = 1.00$) and trial-bin ($F(3, 21) = 3.55$, $p = .032$, $\eta_p^2 = .34$). The interactions were non-significant for any combinations of the factors ($Fs \leq 1.11$, $p \geq .33$, $\eta_p^2 \leq .14$).

A three-way repeated-measures ANOVA (2 stimulus sides × 10 $T_S$ × 4 trial-bins) of $\bar{T}_R$ values in Experiment S2 showed no significant main effect of stimulus side ($F(1, 7) = .037$, $p = .85$, $\eta_p^2 = .0052$) and trial-bin ($F(3, 21) = 1.21$, $p = .33$, $\eta_p^2 = .15$), although there was a significant main effect of $T_S$ ($F(9, 63) = 1117.64$, $p < .001$, $\eta_p^2 = .99$). The interactions were non-significant for any combinations of factors ($Fs \leq 1.31$, $p \geq .15$, $\eta_p^2 \leq .16$).

### Supplementary Results

#### *Experiment S1*

In Experiment S1, $\bar{T}_R(\mu_{\text{2 priors}})$ and $\hat{\mu}_{\text{prior}}$ values were plotted higher than 1059 ms ($\mu_{\text{prior}}$) in trials 1–160 but gradually approached 1059 ms along the progression of trials (Fig. S1a, b). A one-way repeated-measures ANOVA of the $\bar{T}_R(\mu_{\text{2 priors}})$ values exhibited a significant main effect of trial bin ($F(3, 21) = 5.45$, $p = .0062$, $\eta_p^2 = .44$). The results suggested that in the early trials, the participants acquired a higher $\hat{\mu}_{\text{prior}}$ than the actual mean of the wide prior (1059 ms), although they gradually learned the actual mean as $\hat{\mu}_{\text{prior}}$.

#### *Experiments S2*

In Experiment S2, $\bar{T}_R(\mu_{2\text{ priors}})$ and $\hat{\mu}_{\text{prior}}$ values differed between the short and long priors from early trials and had little change over trials (Fig. S1c, d). The $\bar{T}_R(\mu_{2\text{ priors}})$ values were significantly greater for the long prior than for the short prior over all trial bins ($ps_{cor} \leq .0040$ corrected by the Holm method, $ts(7) \geq 3.67$, Cohen's $ds \geq 1.30$), indicating that the participants quickly learned the short and long priors.

In addition, a two-way repeated-measures ANOVA (2 priors × 4 trial-bins) of the $\bar{T}_R(\mu_{2\text{ priors}})$ values in Experiment S2 exhibited a significant main effect of prior ($F(1, 7) = 66.99$, $p < .001$, $\eta_p{}^2 = .91$) but revealed a non-significant main effect of trial bin ($F(3, 21) = .092$, $p = .96$, $\eta_p{}^2 = .013$) and non-significant interaction between them ($F(3, 21) = 2.47$, $p = .090$, $\eta_p{}^2 = .26$). These results indicated that the participants quickly completed learning the priors.

For Experiment S2, we also calculated the regression indices for the short and long priors similar to Experiments 1–5. In all trial bins, the indices were greater than zero for both priors ($ps_{cor} \leq .0070$, $ts(7) \geq 3.25$, $ds \geq 1.15$), and greater for the long prior than for the short prior ($ps_{cor} = .0020$, $ts(7) \geq 4.20$, $ds \geq 1.49$). The results further supported that the participants quickly learned the short and long priors, respectively.

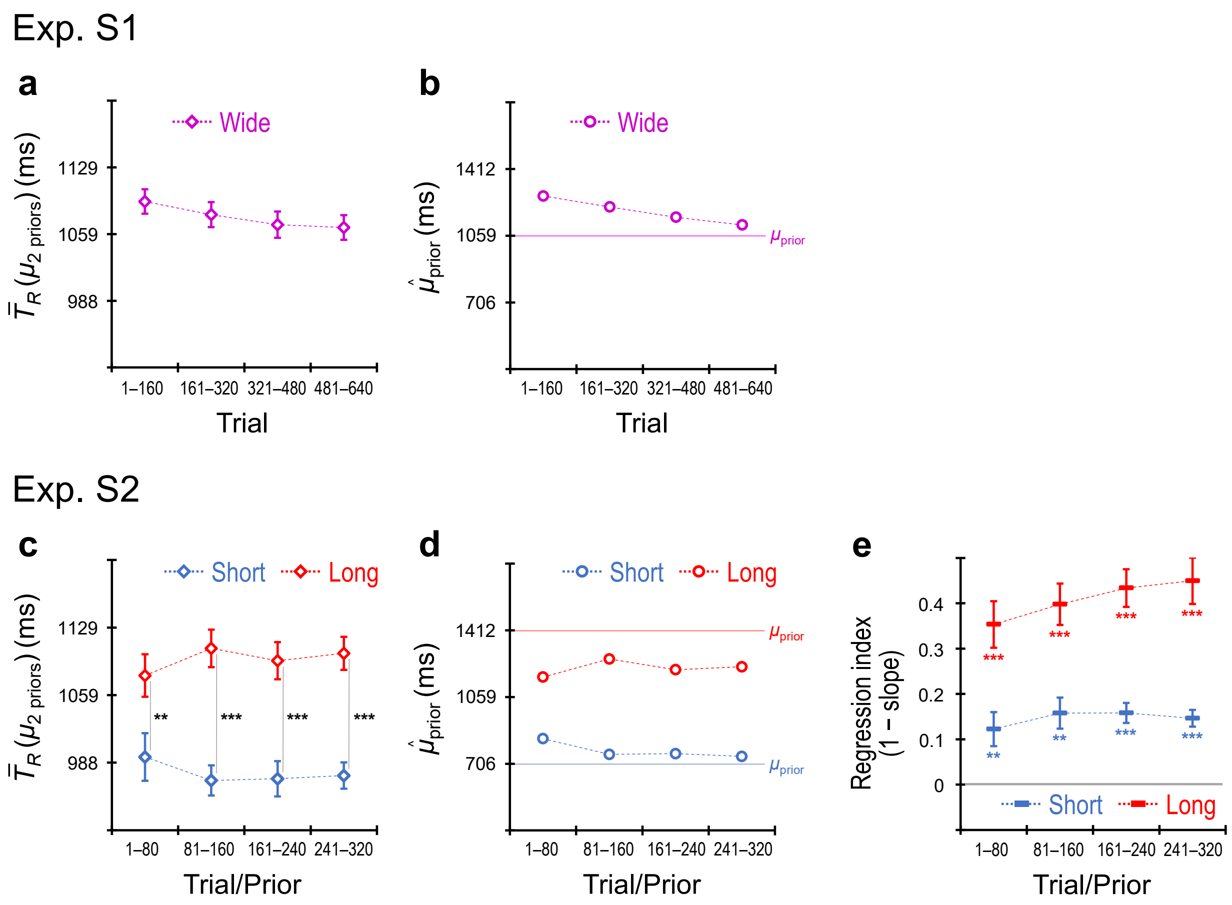


**Supplementary Fig S1. Results for Experiments S1 and S2.** $\bar{T}_R(\mu_{2\ \mathrm{priors}})$ values across the participants (mean ± SEM) for the wide prior in Experiment S1, calculated per 160 trials (**a**). $\hat{\mu}_{\mathrm{prior}}$ values detected using the grand-averaged $\bar{T}_R$ values for the wide prior in Experiment S1, calculated per 160 trials (**b**). $\bar{T}_R(\mu_{2\ \mathrm{priors}})$ values across the participants for the short and long priors in Experiment S2, calculated per 80 trials/prior (**c**). $\hat{\mu}_{\mathrm{prior}}$ values detected using the grand-averaged $\bar{T}_R$ values for the short and long priors in Experiment S2, calculated per 80 trials/prior (**d**). Regression indices across the participants (mean ± SEM) for the short and long priors in Experiment S2, calculated per 80 trials/prior (**e**). ** $p_{cor} < .01$, *** $p_{cor} < .001$.

#### *Variables obtained by curve fittings in Experiments 1–5, S1, and S2*

Tables S2–S6 show the $\hat{\mu}_{\text{prior}}$, $\hat{\sigma}_{\text{prior}}$, and *w* values obtained by the curves fitted to the grand-averaged $\overline{T}_R$ values in Experiments 1–5, S1, and S2. These tables are presented in conjunction with the indication of the values to account for the discrepancy between the results using the $\overline{T}_R(\mu_{2\text{ priors}})$ values and regression indices in trials 1–160 of Experiment 3 (see Experiment 3 in Results and Fig. 6b and d).

Notably, the $\hat{\mu}_{\text{prior}}$ ($\mu_{\text{prior}}$ in Eq. 4) is uniquely specified from the fitted curves based on Eq. 4, whereas $\hat{\sigma}_{\text{prior}}$ ($\sigma_{\text{prior}}$ in Eq. 4) and *w* are unspecified and vary depend on the initial values used for the fittings. To calculate the $\hat{\sigma}_{\text{prior}}$ and *w* values shown in the tables, two types of initial values were used as follows: (1) $\hat{\mu}_{\text{prior}}$ (short/long) = 1059 ms, $\hat{\sigma}_{\text{prior}}$ (short/long) = 427 ms, *w* = 0.15 (Supplementary Tables S3 and S5), (2) $\hat{\mu}_{\text{prior}}$ (short) = 706 ms, $\hat{\mu}_{\text{prior}}$ (long) = 1412 ms, $\hat{\sigma}_{\text{prior}}$ (short/long) = 223 ms, *w* = 0.15 (Supplementary Tables S4 and S6). The initial values of $\hat{\mu}_{\text{prior}}$ and $\hat{\sigma}_{\text{prior}}$ in type (1) were based on the physical statics of the generalised or wide prior; those in type (2) were based on the physical statics of the short and long priors. The initial value of *w* = 0.15 was commonly used in both types, which was based on a previous study [12].

**Supplementary Table S2. The $\hat{\mu}_{\text{prior}}$ values (ms) obtained by fitting the curves based on Eq. 4 to the grand-averaged $\overline{T}_R$ values as a function of $T_S$**

| Exp. # | Prior | Trial bin | | | |
|---|---|---|---|---|---|
| | | **1–160** (1–80/Prior) | **161–320** (81–160/Prior) | **321–480** (161–240/Prior) | **481–640** (241–320/Prior) |
| 1 | Short | 1226.2 | 1242.1 | 1315.0 | 1349.6 |
| | Long | 1234.5 | 1280.8 | 1273.8 | 1225.4 |
| 2 | Short | 1169.5 | 985.4 | 942.7 | 683.4 |
| | Long | 1242.3 | 1249.7 | 1190.3 | 1206.6 |
| 3 | Short | **1158.1** | 1014.2 | 933.9 | 941.0 |
| | Long | 1370.9 | 1364.0 | 1358.6 | 1299.2 |
| 4 | Short | 1060.5 | 1061.4 | 1007.7 | 988.1 |
| | Long | 1389.5 | 1409.6 | 1382.0 | 1395.0 |
| 5 | Short | 1021.5 | 1016.6 | 1012.7 | 890.8 |
| | Long | 1320.8 | 1384.5 | 1356.8 | 1337.9 |
| S1 | Wide | 1269.4 | 1211.2 | 1156.8 | 1116.1 |
| S2 | Short | 838.6 | 755.6 | 759.4 | 744.7 |
| | Long | 1164.5 | 1260.6 | 1203.5 | 1219.6 |

These $\hat{\mu}_{\text{prior}}$ values were obtained regardless of whether using the initial values (1) or (2), except for the minute difference below significant digits. The bold value (**1158.1** ms) indicates the $\hat{\mu}_{\text{prior}}$ value for the short priors in trials 1–160 of Experiment 3 (*cf.* the last paragraph of 'Experiment 3: timing using two body parts (right vs left hands)' in Results).

**Supplementary Table S3. The $\hat{\sigma}_{\text{prior}}$ values (ms) obtained by fitting the curves based on Eq. 4 to the grand-averaged $\overline{T}_R$ values as a function of $T_S$**

| Exp. # | Prior | Trial bin | | | |
|---|---|---|---|---|---|
| | | 1–160 (1–80/Prior) | 161–320 (81–160/Prior) | 321–480 (161–240/Prior) | 481–640 (241–320/Prior) |
| 1 | Short | 415.8 | 419.8 | 460.4 | 470.8 |
| | Long | 382.2 | 390.4 | 380.7 | 398.6 |
| 2 | Short | 381.9 | 384.8 | 422.5 | 505.3 |
| | Long | 363.5 | 378.9 | 398.3 | 371.9 |
| 3 | Short | **365.3** | 393.4 | 367.5 | 431.1 |
| | Long | 360.2 | 389.2 | 332.3 | 390.4 |
| 4 | Short | 334.0 | 334.2 | 342.9 | 336.8 |
| | Long | 375.2 | 381.7 | 395.8 | 384.6 |
| 5 | Short | 351.6 | 358.9 | 383.0 | 347.6 |
| | Long | 383.9 | 377.1 | 372.7 | 359.9 |
| S1 | Wide | 378.4 | 413.7 | 450.0 | 457.6 |
| S2 | Short | 339.8 | 350.7 | 341.3 | 376.3 |
| | Long | 397.1 | 374.5 | 368.0 | 360.2 |

Type (1) of the initial values were used for fitting. The bold value (**365.3** ms) indicates the $\hat{\sigma}_{\text{prior}}$ value for the short priors in trials 1–160 of Experiment 3 (*cf.* the last paragraph of 'Experiment 3: timing using two body parts (right vs left hands)' in Results).

**Supplementary Table S4. The $\hat{\sigma}_{\text{prior}}$ values (ms) obtained by fitting the curves based on Eq. 4 to the grand-averaged $\overline{T}_R$ values as a function of $T_S$**

| Exp. # | Prior | Trial bin | | | |
|---|---|---|---|---|---|
| | | **1–160** (1–80/Prior) | **161–320** (81–160/Prior) | **321–480** (161–240/Prior) | **481–640** (241–320/Prior) |
| 1 | Short | 296.6 | 299.3 | 381.2 | 385.6 |
| | Long | 341.0 | 312.8 | 306.0 | 304.4 |
| 2 | Short | 270.5 | 283.8 | 349.6 | 436.7 |
| | Long | 284.3 | 279.7 | 290.1 | 291.0 |
| 3 | Short | **256.9** | 294.2 | 300.3 | 358.4 |
| | Long | 284.7 | 311.3 | 311.1 | 257.3 |
| 4 | Short | 226.7 | 226.7 | 233.0 | 226.0 |
| | Long | 297.3 | 303.3 | 313.1 | 307.2 |
| 5 | Short | 242.2 | 248.3 | 281.3 | 229.2 |
| | Long | 308.6 | 298.3 | 298.0 | 287.2 |
| S1 | Wide | 313.8 | 341.5 | 376.2 | 360.3 |
| S2 | Short | 243.4 | 248.6 | 240.8 | 254.2 |
| | Long | 288.8 | 264.9 | 259.2 | 251.5 |

Type (2) of the initial values were used for fittings. The bold value (**256.9** ms) indicates the $\hat{\sigma}_{\text{prior}}$ value for the short priors in trials 1–160 of Experiment 3 (*cf.* the last paragraph of 'Experiment 3: timing using two body parts (right vs left hands)' in Results).

**Supplementary Table S5. The *w* values obtained by fitting the curves based on Eq. 4 to the grand-averaged $\overline{T}_R$ values as a function of $T_S$**

| Exp. # | Prior | Trial bin | | | |
|---|---|---|---|---|---|
| | | **1–160** (1–80/Prior) | **161–320** (81–160/Prior) | **321–480** (161–240/Prior) | **481–640** (241–320/Prior) |
| 1 | Short | 0.157 | 0.154 | 0.123 | 0.118 |
| | Long | 0.163 | 0.163 | 0.165 | 0.159 |
| 2 | Short | 0.181 | 0.185 | 0.152 | 0.126 |
| | Long | 0.166 | 0.179 | 0.174 | 0.163 |
| 3 | Short | **0.194** | 0.177 | 0.165 | 0.149 |
| | Long | 0.175 | 0.166 | 0.141 | 0.169 |
| 4 | Short | 0.218 | 0.219 | 0.215 | 0.221 |
| | Long | 0.171 | 0.169 | 0.163 | 0.168 |
| 5 | Short | 0.206 | 0.202 | 0.186 | 0.226 |
| | Long | 0.166 | 0.171 | 0.170 | 0.173 |
| S1 | Wide | 0.160 | 0.154 | 0.144 | 0.157 |
| S2 | Short | 0.203 | 0.204 | 0.207 | 0.210 |
| | Long | 0.174 | 0.194 | 0.196 | 0.203 |

Type (1) of the initial values were used for fittings. The bold value (**0.194**) indicates the *w* value for the short priors in trials 1–160 of Experiment 3 (*cf*. the last paragraph of ‘Experiment 3: timing using two body parts (right vs left hands)’ in Results).

**Supplementary Table S6. The *w* values obtained by fitting the curves based on Eq. 4 to the grand-averaged $\overline{T}_R$ values as a function of $T_S$**

| Exp. # | Prior | Trial bin 1–160 (1–80/Prior) | 161–320 (81–160/Prior) | 321–480 (161–240/Prior) | 481–640 (241–320/Prior) |
|---|---|---|---|---|---|
| 1 | Short | 0.112 | 0.110 | 0.101 | 0.097 |
| | Long | 0.145 | 0.130 | 0.133 | 0.121 |
| 2 | Short | 0.128 | 0.136 | 0.126 | 0.109 |
| | Long | 0.130 | 0.132 | 0.126 | 0.127 |
| 3 | Short | **0.136** | 0.133 | 0.135 | 0.124 |
| | Long | 0.138 | 0.133 | 0.132 | 0.111 |
| 4 | Short | 0.148 | 0.148 | 0.146 | 0.149 |
| | Long | 0.136 | 0.135 | 0.129 | 0.134 |
| 5 | Short | 0.142 | 0.140 | 0.137 | 0.149 |
| | Long | 0.133 | 0.135 | 0.136 | 0.138 |
| S1 | Wide | 0.132 | 0.127 | 0.120 | 0.123 |
| S2 | Short | 0.145 | 0.145 | 0.146 | 0.142 |
| | Long | 0.127 | 0.137 | 0.138 | 0.142 |

Type (2) of the initial values were used for fittings. The bold value (**0.136**) indicates the *w* value for the short priors in trials 1–160 of Experiment 3 (*cf*. the last paragraph of 'Experiment 3: timing using two body parts (right vs left hands)' in Results).
